# Metabolic analysis of sarcopenic muscle identifies positive modulators of longevity and healthspan in *C. elegans*

**DOI:** 10.1101/2024.10.19.616468

**Authors:** Steffi M Jonk, Alan Nicol, Vicki Chrysostomou, Emma Lardner, Shu-Che Yu, Gustav Stålhammar, Jonathan G Crowston, James R Tribble, Peter Swoboda, Pete A Williams

## Abstract

Sarcopenia is the age-related degeneration of skeletal muscle, resulting in loss of skeletal muscle tone, mass, and quality. Skeletal muscle is a source of systemic metabolites and macromolecules important for neuronal health, function and healthy neuronal aging. Age-related loss of skeletal muscle might result in decreased metabolite and macromolecule availability, resulting in reduced neuronal function or increased susceptibility to unhealthy aging and neurodegenerative diseases. We aimed to identify muscle metabolite candidates that regulate healthy aging. C57BL/6J mice were aged to young adult (4 months) and old age (25 months) and skeletal muscle was collected. Age related muscle loss was confirmed by reduced muscle mass, muscle fiber degeneration, reduced myosin intensity, in addition to a metabolic shift and increased DNA damage in skeletal muscle. Using a low molecular weight enriched metabolomics protocol, we assessed the metabolic profile of skeletal muscle from young adult and old mice and identified 20 metabolites that were significantly changed in aged muscle. These candidate metabolites were tested in *C. elegans* assays of lifespan, health span, muscle-, and mitochondrial morphology under normal and stressed conditions. We identified four candidate metabolites (beta-alanine, 4-guanidinobutanoic acid, 4-hydroxyproline, pantothenic acid) that when supplemented in *C. elegans* provided robust gero- and mitochondrial protection. These candidates also affected life-, and health span in *C. elegans* models of amyotrophic lateral sclerosis and Duchenne muscular dystrophy. Our findings support that aging muscle can be used to identify novel metabolite modulators of lifespan and health and may show promise for future treatments of neurodegenerative and neuromuscular disorders.

## Introduction

Sarcopenia is the age-related degeneration of skeletal muscle characterized by the progressive degeneration of muscle weight, tone and/or mass, resulting in reduced muscle capacity and strength. The prevalence of sarcopenia is 5-13% for humans aged 60–70 and 11-50% for those aged >80 years^1, 2^. In sarcopenia, myofiber atrophy and a loss of myofiber numbers occurs, in addition to a loss of myofiber function due to neuromuscular junction destabilization^3^. Inflammation is a key hallmark of aging, as well as sarcopenia, and inhibition and deletion of a pro-inflammatory cytokine (IL-11) improved metabolic function and muscle quality in mice^4^. In aged muscle and sarcopenia, there is a greater loss of type II (fast twitch) muscle fibers than type I (slow twitch), leading to an increased ratio of type I fibers and a respiratory switch, which exerts increased relaxation time and a loss of power^5, 6^. This loss of power contributes to accidents, disability, and a poor quality of life in addition to correlating with cardiac and respiratory disease, cognitive impairment, and neurodegenerative disease^7^.

Skeletal muscle is an important source and store of amino acids and regulator of metabolism by responding dynamically to bioenergetic demand^1^. To achieve these dynamic roles, muscle secretes amino acids, proteins, lipids, metabolites, cytokines (myokines), and small RNAs; collectively referred to as the muscle secretome^8^. Remaining physically active and high intensity training is not only an intervention for sarcopenia, but also protects against neurodegenerative diseases. Supporting this, forced and voluntary exercise is profoundly neuroprotective in many animal models of neurodegenerative disease^9, 10, 11, 12, 13^. Together, this data supports the importance of systemic and active secretion of macromolecules and metabolites by skeletal muscle. Supplementation of macromolecules and metabolites and their effect on disease, health span and lifespan have therefore been an ongoing research topic and intervention strategy^14, 15, 16^.

Here, we demonstrate changes in the muscle metabolome during aging. The muscle metabolome of aged mice was used to identify altered metabolites with age and sarcopenia. Supplementation of these candidate metabolites in *C. elegans* extended lifespan, improved health span, and affected mitochondrial and myosin degeneration. The identified metabolite candidates also provided therapeutic benefit in *C. elegans* models of neuromuscular disease (amyotrophic lateral sclerosis; ALS, and Duchenne muscular dystrophy; DMD). This study provides new strategies for gero- and metabolic-protection by targeting key metabolic pathways.

## Results and Discussion

The age-related, or sarcopenic, loss of muscle mass is well-reported in human patients and model organisms^17^. However, there are clinically defined standards to diagnose sarcopenia in humans, but not in mice models, and it is therefore standard to analyse body and muscle weight in mice to define age related muscle loss, which may reflect sarcopenia in mice^18^. In a recent natural history study of sarcopenia, the body mass of male C57BL/6JRj mice was identified stable between 8-18 months, before it started to decline, and mice had lost ∼10% of their body mass at 24 months of age^1^. To assess the metabolic profile of aged muscle we aged C57BL/6J (B6J) mice to 25 months (representing ∼70+ years of age in humans)^19^. At 25 months, B6J mice had unchanged body weight (**Fig. 1A**), whilst the weight and overall area of the skeletal muscles gastrocnemius and quadriceps decreased (**Fig. 1B, C**), with a concomitant muscle fibre degeneration (**Fig. 1D-F**). While the muscle degenerates, fat cells (adipocytes) infiltrate the muscle tissue, which associates to muscle dysfunction in disease- and aging states^20,21^. We identified adipocyte infiltration in skeletal muscle, which was more prevalent in the quadriceps than in the gastrocnemius (**Fig. S1A, B**). The gastrocnemius and quadriceps are skeletal muscles, located in the lower leg and front of the thigh, and are characterized by a high proportion of type II, fast-twitch fibres that have high glycolytic capacity to rapidly generate ATP^22^. During aging and in sarcopenia, both type I and type II fibres degenerate but with a preference for type II fibre degeneration^23^. Fibre types are generally characterized by the expression of myosin heavy chain (MyHC) isotypes^24^, and we confirmed sarcopenia, by a decrease of myosin, representing both type I and II fibres, whilst MYH7B increased (type I fibre specific myosin (**Fig. 1G, H**), suggesting an increase in the proportion of type I fibres.

**Figure. 1.**
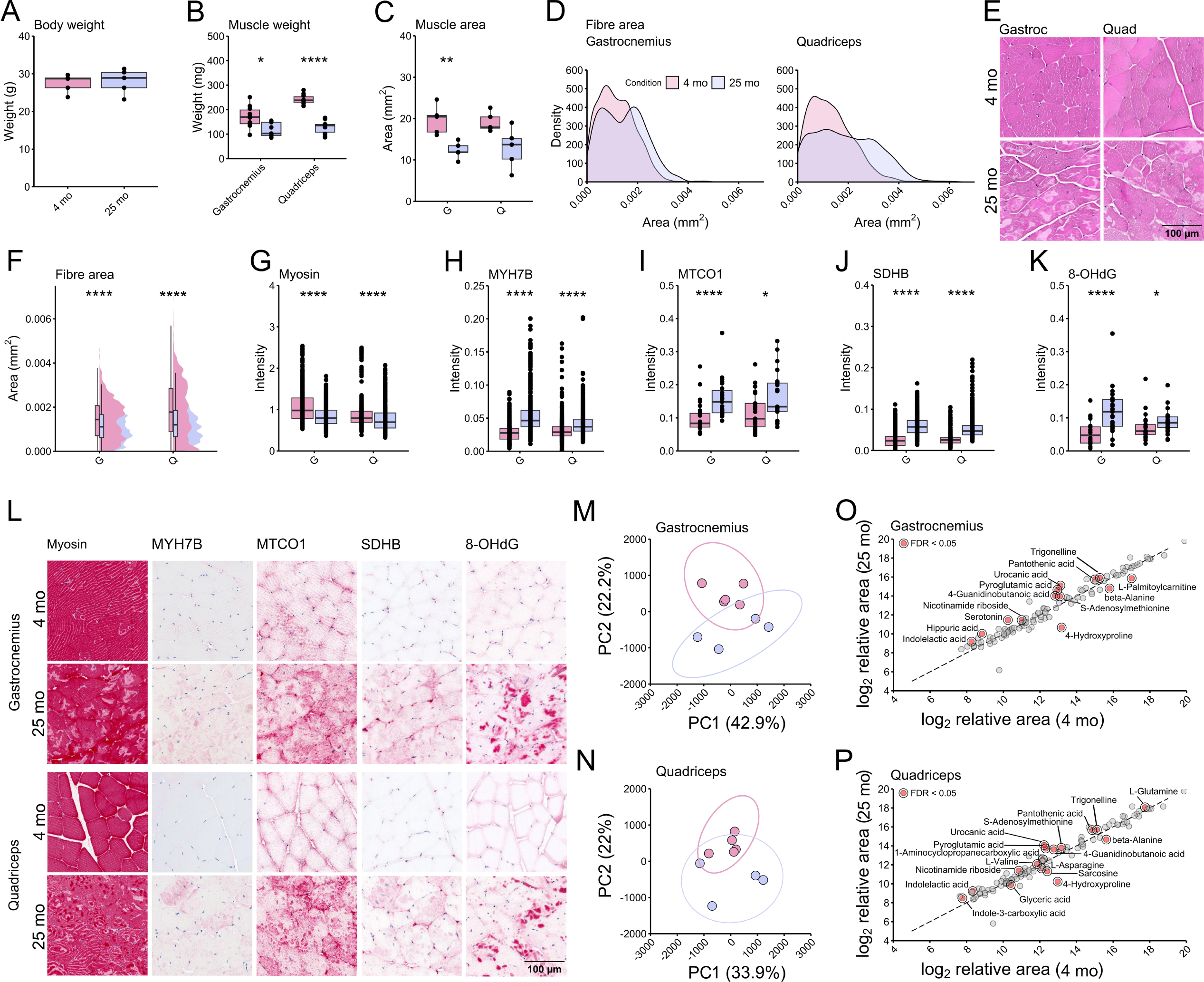
Metabolic changes in aging murine skeletal muscle. (**A-P**) C57BL/6J mice. (**A**) Body weight did not change during aging (n=5/group). (**B**) Muscle weight and (**C**) area declined during aging, as did (**D-F**) fiber area, confirming sarcopenia. By histological assessment (**F-L**), it was observed that (**G**) myosin intensity declined while (**H**) MHY7B intensity increased, suggesting an increase in the ratio of type I (slow twitch) fibers. In line with this finding, (**I**) MTCO1, a component of cytochrome c oxidase, increased, together with (**J**) SDHB, a subunit of succinate dehydrogenase, suggesting a respiratory switch from glycolytic to oxidative. (**K**) 8-OHdG intensity increased, identifying cell death in aged tissue. (**M-N**) Unsupervised principal component analysis (PCA) separated samples into discrete groups that confirmed distinct metabolic profiles of young adult and aged muscle tissue. Statistical analysis of the low molecular weight metabolome identified 17 metabolites in (**O**) gastrocnemius and 13 in (**P**) quadriceps (significant hits in red). 10 were identical between the datasets, leaving 20 unique metabolites (t-test FDR <0.05, volcano plot FDR <0.1). G: gastrocnemius, Q: quadriceps. *Pink* = 4 months of age, *purple* = 25 months of age. *p = * < 0.05, ** < 0.01, *** < 0.001, **** < 0.0001*.

Mitochondrial density and function is of great importance in skeletal muscle, generating ATP rapidly at sites of action and accumulating at neuromuscular junctions (NMJs)^25, 26^. Mitochondrial degeneration is a hallmark of sarcopenia and a target for treatment strategies in sarcopenia^27^. We next assessed gross metabolic potential using histopathological techniques (**Fig. 1L**) that demonstrated an increase in MTCO1 intensity (cytochrome *c* oxidase; mitochondrial Complex IV) (**Fig. 1I**). No differences were observed for CYOC (Cytochrome C Oxidase subunit Vic/COX6C), another component of cytochrome *c* oxidase (**Fig. S1A, C**). Cytochrome *c* oxidase is a biomarker for mitochondrial injury, oxidative stress and ROS production, and apoptosis and is often used to diagnose mitochondrial disease^28, 29^. Succinate dehydrogenase complex iron sulfur subunit B (SDHB), a protein subunit of succinate dehydrogenase (mitochondrial Complex II), increased during aging, supporting an increase of oxidative respiration due to an increase of the proportion of type I fibres (**Fig. 1J**). DNA damage is a common aging hallmark^30^. DNA damage was characterized by the presence of 8-OHdG, targeting hydroxy deoxyguanosine, a modified base in DNA due to attack by hydroxyl radicals (formed as byproducts of aerobic metabolism and ROS), demonstrating an increase of DNA damage in aged muscle tissue (**Fig. 1K**). Collectively, this data demonstrates aged tissue and metabolic compromise in the skeletal muscle of 25-month-old mice.

We next conducted metabolomic profiling in young and aged muscle. We utilized a low molecular weight metabolomics platform to measure 502 low molecular weight metabolites in skeletal muscle^31^. The resulting skeletal muscle metabolomes demonstrated a clear change with advancing age. Principal component analysis confirmed distinct metabolic profiles of young adult and aged muscle tissue but showing variation between samples by PC1 (principal component 1) and PC2 (**Fig. 1M, N**). A total of 30 metabolites were identified as significantly changed in gastrocnemius and quadriceps, of which 10 were identical between the datasets, leaving 20 unique metabolites (**Fig. 1O, P**, **Table 1**). KEGG (Kyoto Encyclopedia of Genes and Genomes) enrichment analysis identified *arginine and proline*-, *beta-alanine-, histidine-, glycine, serine and threonine-, glyoxylate and dicarboxylate-, nicotinate*, and *nicotinamide metabolism*, and *pantothenate and CoA biosynthesis* as the most affected pathways. P*antothenate and CoA biosynthesis* were identified as significant, leaving the other pathways as matched with the identified metabolites but not predicted to be significantly changed (**Fig. S1D**). RaMP DB (Relation database of Metabolic Pathways) enrichment identified several transport processes as neurotransmitter transporters, transport of small molecules, transmembrane transport, and metabolic processes as histidine catabolism and metabolism, metabolism of amino acids and derivatives, and metabolic disorders of biological oxidation enzymes (**Fig. S1E**).

**Table 1.**
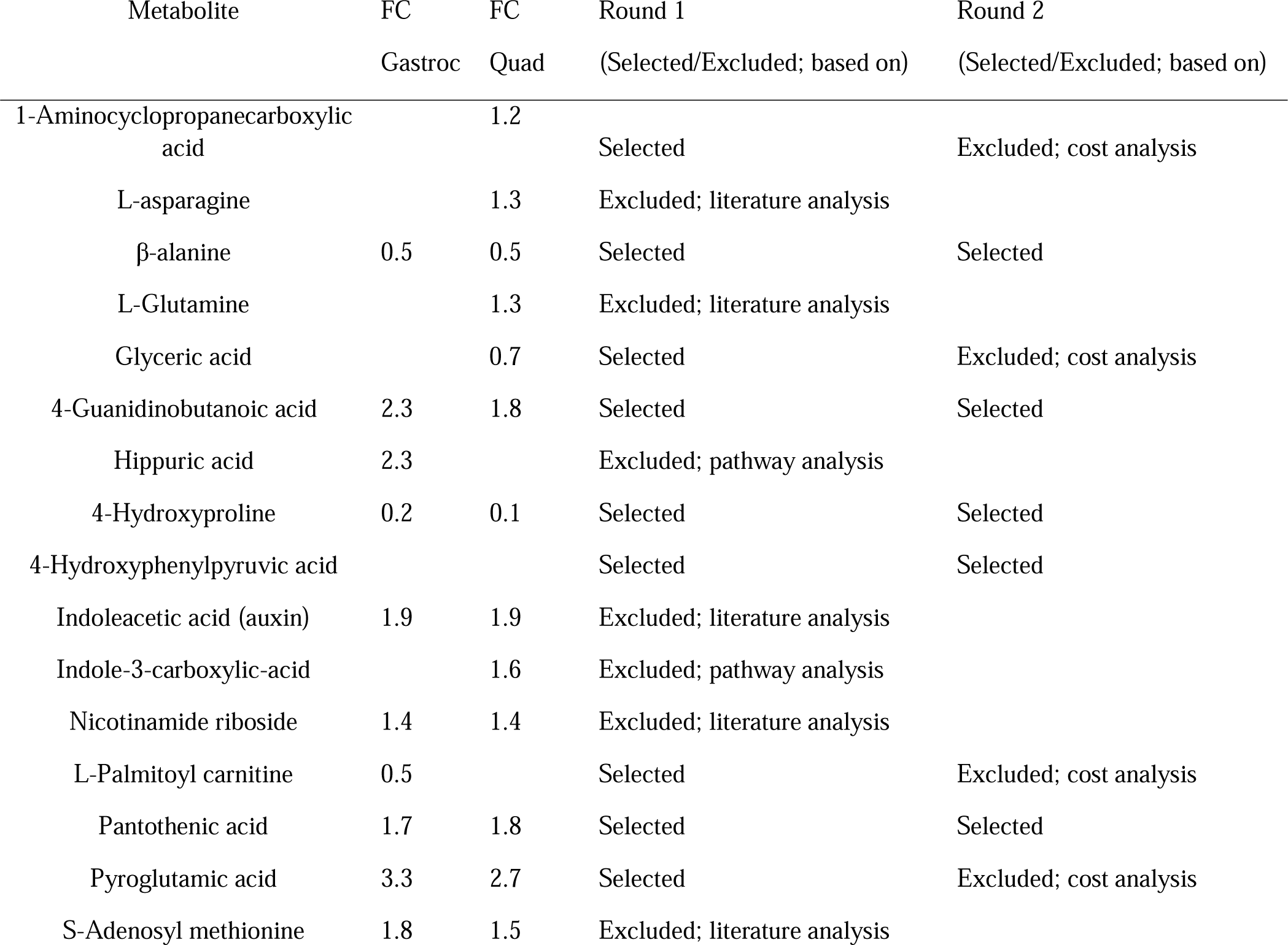
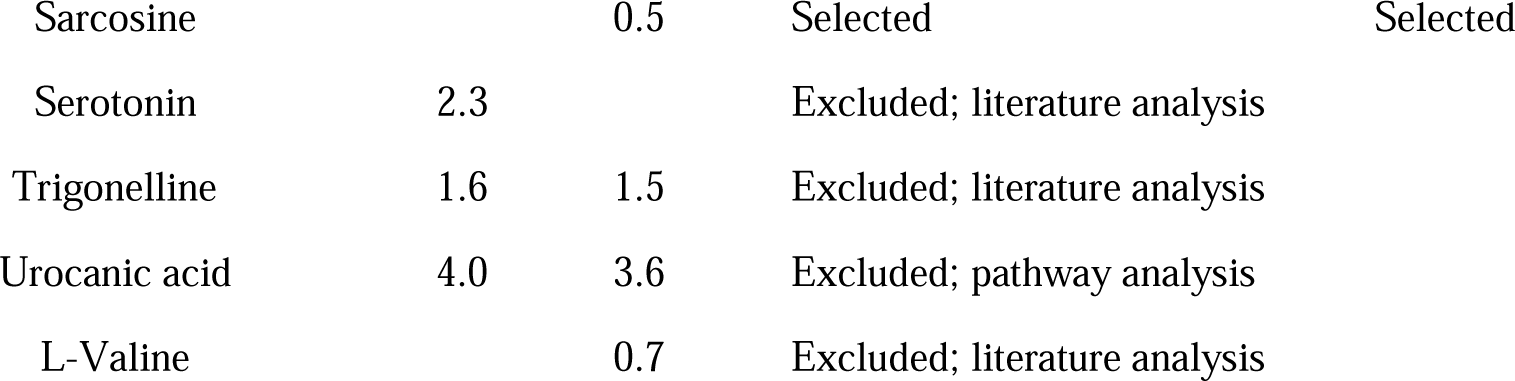
Significant hits and triaging of metabolites. Metabolites identified in the low molecular weight metabolomics dataset were triaged for follow-up studies in *C. elegans* assays. Triaging was performed by literature- and pathway analysis for each significant hit (p-value and fold-change, 4-hydroxyphenylpyruvic acid was not identified by volcano plot but was included based on FC). Literature analysis was performed in PubMed and Textpresso (search strategy by metabolite + mice/*C. elegans*/lifespan), and pathways analysis in *C. elegans (*KEGG and wormflux) for each metabolite^50, 51^. Metabolites that were identified as excretory products were labelled as not relevant. Based on the literature analysis, eight metabolites were excluded since (extensive) literature related to longevity in mice and/or *C. elegans* was published; lifespan effects of metabolite candidates were well studied and published. Based on pathway analysis, two candidates were labelled as excretory products and excluded, one was not identified in known pathways (KEGG). The remaining novel and relevant candidates were further triaged based on availability and cost. All candidates were commercially available; however, 4 candidates were excluded based on a cost analysis and experiments not being feasible. FC = fold change old/young.

| Metabolite | FC | FC | Round 1 | Round 2 |
| --- | --- | --- | --- | --- |
|  | Gastroc | Quad | (Selected/Excluded; based on) | (Selected/Excluded; based on) |
| 1-Aminocyclopropanecarboxylic acid |  | 1.2 | Selected | Excluded; cost analysis |
| L-asparagine |  | 1.3 | Excluded; literature analysis |  |
| β-alanine | 0.5 | 0.5 | Selected | Selected |
| L-Glutamine |  | 1.3 | Excluded; literature analysis |  |
| Glyceric acid |  | 0.7 | Selected | Excluded; cost analysis |
| 4-Guanidinobutanoic acid | 2.3 | 1.8 | Selected | Selected |
| Hippuric acid | 2.3 |  | Excluded; pathway analysis |  |
| 4-Hydroxyproline | 0.2 | 0.1 | Selected | Selected |
| 4-Hydroxyphenylpyruvic acid |  |  | Selected | Selected |
| Indoleacetic acid (auxin) | 1.9 | 1.9 | Excluded; literature analysis |  |
| Indole-3-carboxylic-acid |  | 1.6 | Excluded; pathway analysis |  |
| Nicotinamide riboside | 1.4 | 1.4 | Excluded; literature analysis |  |
| L-Palmitoyl carnitine | 0.5 |  | Selected | Excluded; cost analysis |
| Pantothenic acid | 1.7 | 1.8 | Selected | Selected |
| Pyroglutamic acid | 3.3 | 2.7 | Selected | Excluded; cost analysis |
| S-Adenosyl methionine | 1.8 | 1.5 | Excluded; literature analysis |  |
| Sarcosine |  | 0.5 | Selected | Selected |
| Serotonin | 2.3 |  | Excluded; literature analysis |  |
| Trigonelline | 1.6 | 1.5 | Excluded; literature analysis |  |
| Urocanic acid | 4.0 | 3.6 | Excluded; pathway analysis |  |
| L-Valine |  | 0.7 | Excluded; literature analysis |  |

We triaged 20 metabolite candidates based on existing literature and pathway analysis, for lifespan assessment in *C. elegans,* since this is a short-lived model organism that is widely used for lifespan analysis^32, 33^. The candidates were then investigated in *C. elegans* (triaging strategy at **Table 1**); where we performed a flooding assay (*C. elegans* were flooded with M9 buffer to assess through thrashing live/dead worms) at days 12, 16, and 20 to initially screen for potential lifespan enhancers (**Fig. S2A, Table S4**). Wild-type N2 *C. elegans* are young adults at day 2, start degenerating around day 8, reach old age around day 12 and very old age beyond day 16 (this is matched in our laboratory)^34,35^. Based on this data, long-term survival assays were performed for the six top candidate metabolites with lifespan enhancing potential: β-alanine, glycine, 4-hydroxyphenylpyruvic acid (4HPPA), 4-hydroxyproline, 4-guanidinobutanoic acid, and pantothenic acid. Supporting our triaging strategy, glycine extended *C. elegans* lifespan as reported previously (**Fig. S2B**) ^36^, as did novel candidates β-alanine (BA), 4-hydroxyproline (4HP), 4-guanidinobutanoic acid (4GBA) and pantothenic acid (PA) (**Table 1**, **Fig. 2A-B, Table S5**).

**Figure. 2.**
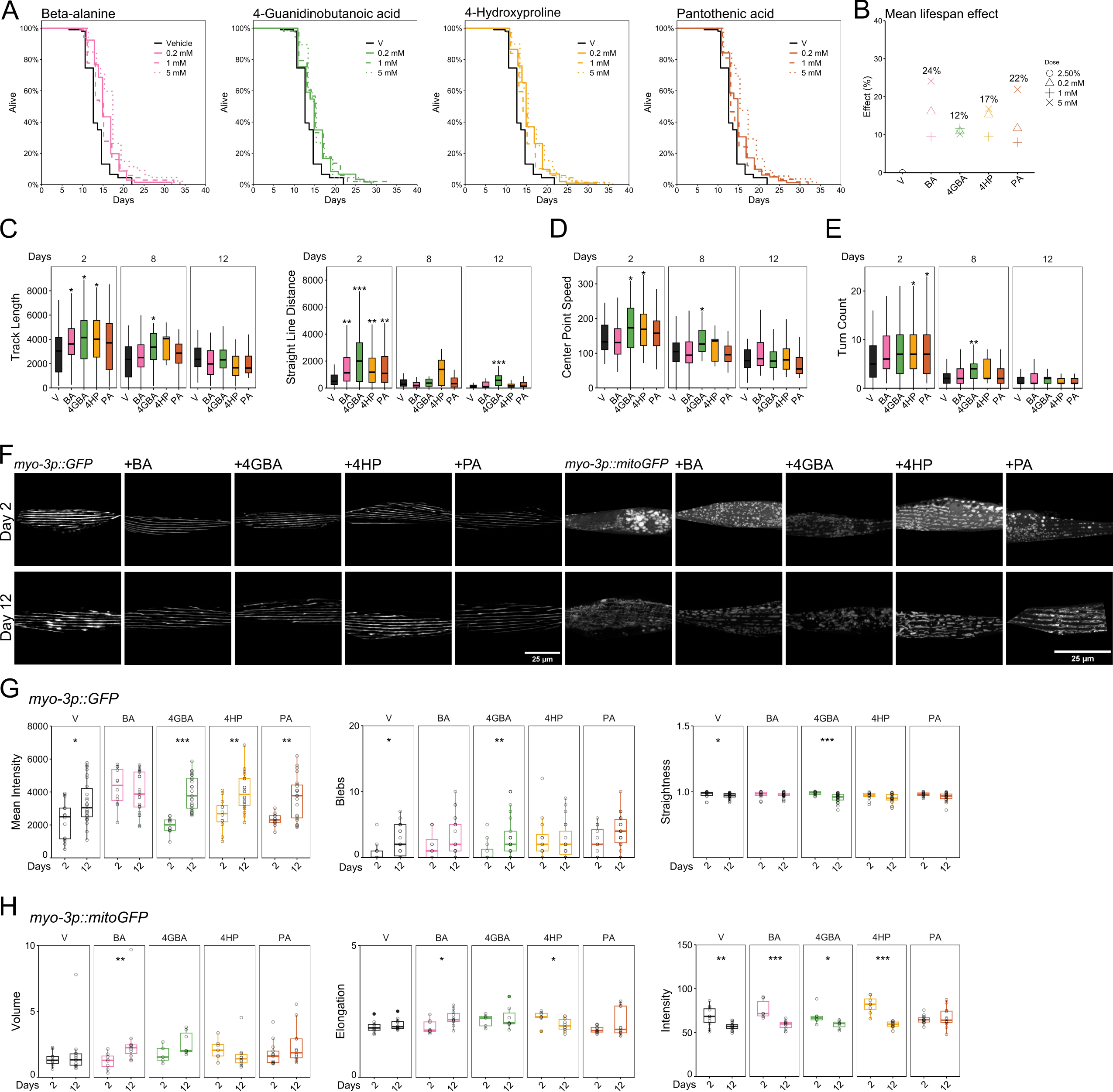
Identification of novel metabolite candidates for gero- and metabolic protection in *C. elegans*. (**A-E**) *C. elegans*, wild-type N2, (**F-G**) *C. elegans*, RW1596, (**F, H**) *C. elegans*, SJ4103. (**A**) Based on literature, pathway, and cost-analysis, we identified 4 top hits, which all increased lifespan, shown in days. (**B**) Based on the survival curve, the mean lifespan effect was identified, which ranged from 10 to 25%. (**C**) Health span by movement track length and straight-line distance (WormLab) increased at day 2 and day 8, but not at day 12. (**D**) Health span by centre point speed and (**E**) turn count (WormLab) increased at day 2 and day 8, but not at day 12. (**F**) Fluorescence microscopy images representing *myo-3p::GFP* at day 2 and day 12, and confocal microscopy images representing *myo-3p::mitoGFP* at day 2 and day 12. During aging, myosin fibres degenerate, lose straightness and increase in the number of blebs. Mitochondrial degeneration is observed by a loss of mean intensity. (**G**) Myosin fibres were quantified by MuscleMetrics, plots represent the mean intensity, number of blebs and straightness for all supplements at day 2 and day 12 (*n = number of sarcomeres:* VH=14/26, BA=12/23, 4GBA=12/25, 4HP=12/20, PA=12/22). (**H**) Mitochondria were quantified by Imaris and MitoAnalyzer, plots represent the volume, elongation, and intensity for all supplements at day 2 and day 12 (*n = number of worms, average of 2 regions per worm:* VH=11/14, BA=6/10, 4GBA=6/7, 4HP=8/11, PA=14/12). V: vehicle, BA: β-alanine, 4GBA: 4-guanidinobutanoic acid, 4HP: 4-hydroxyproline, PA: pantothenic acid*. p = * < 0.05, ** < 0.01, *** < 0.001, **** < 0.0001*.

As BA, 4HP, 4GBA, and PA significantly extended *C. elegans* lifespan, we next assessed health span to test for improved quality of life during aging when supplemented with these metabolite candidates. *C. elegans* health span measurements include preservation of locomotion, feeding behaviour and accumulation of auto-fluorescent pigment. It has previously been reported that short-lived *C. elegans* mutants score worse on locomotion parameters like speed and amplitude^37^. A consensus on parameters to determine health span in *C. elegans* is lacking. However, it was recently suggested that health span can broadly be determined by measuring physiological parameters like oxidative stress resistance, heat stress resistance, thrashing, and distance travelled^38^. We supplemented the metabolite candidates to *C. elegans* during a 20-day period (starting at day 1 of adulthood), which demonstrated that supplemented worms significantly increased locomotion at day 2 of adulthood. Although small increases of activity were noted, none of the supplemented metabolite candidates caused consistently significant effects (at day 8, 12, 16 and 20 of adulthood) on all parameters (**Fig. 2C-E, Fig S2**). Of note, 4GBA exerted a significant increase on track length and straight-line distance at day 2, 8 and 12 of adulthood, which suggests improved locomotion during aging (**Fig. 2C-E**).

Mitochondrial degeneration within skeletal muscle is one of the first hallmarks of sarcopenia. We studied the effect of BA, 4HP, 4GBA, and PA on skeletal muscle and their mitochondria using *C. elegans* that express GFP (green fluorescent protein) under the *myo-3p* (myosin heavy chain) gene promotor, or that express GFP only in mitochondria, under the same promotor (**Fig. 2F**). Recent studies have predicted biological age based on sarcopenia in *C. elegans* where ‘delayed onset of sarcopenia served as a biomarker of extended lifespan’ ^39^. Using this toolbox, we identified that BA preserved several parameters (GFP intensity, number of blebs, muscle straightness), suggesting a delayed onset of sarcopenia. 4GBA showed a similar effect on all parameters as the vehicle condition, suggesting an early sarcopenic state at day 12 of adulthood, whilst 4HP and PA scored similar as vehicle on some but not on all parameters (**Fig. 2F,G, S2G**). Mitochondrial networks are tightly regulated and change during aging. Mitochondrial networks can consist of small, fragmented units or of larger elongated networks, and it has previously been reported that mitochondria elongate during nutrient deprivation, protecting them from autophagy^40^. Mitochondrial volume and network elongation increased following BA supplementation, demonstrating a larger, more elongated network. PA preserved mean GFP intensity. 4HP decreased mitochondrial elongation, while increasing the number of surfaces and sphericity, suggesting a network of fragmented, round units (**Fig. 2F,H**, **Fig S2G**). Future experiments are needed to determine the functional state of aged mitochondria after metabolite candidate supplementation.

As a broad health span analysis involves oxidative stress resistance, we supplemented the metabolite candidates in models of oxidative stress. *C. elegans* N2 worms were exposed to rotenone and paraquat, which are both inducers of metabolic/oxidative stress by inhibition of mitochondrial Complex I, redox cycling, and by the production of reaction oxygen species (ROS)^41, 42^. Under metabolite supplementation, BA, 4GBA, and 4HP increased lifespan in the rotenone model, while PA increased survival in the paraquat model, suggesting that metabolite candidates affect different mitochondrial processes (**Fig. 3A-B, Table S6**). Since the metabolites were able to partially overcome significant rotenone induced metabolic stress, we also assessed their effects in connection to more chronic neuromuscular disease. We therefore applied the candidates in *C. elegans* models of ALS and DMD (**Table S2**). To assess these candidates in the context of ALS we used a *sod-1*G85R^C^ mutant and *sod-1*WT^C^ wild-type strain, whereby the *sod-1*G85R^C^ mutant is a single copy ALS SOD1 knock-in model, which exhibits both cholinergic and glutamatergic neuron degeneration with an increased sensitivity to paraquat exposure^43^. Survival assays identified that BA, 4GBA, 4HP and PA slightly increased mean lifespan of the *sod-1*WT^C^ wildtype strain, but not in the *sod-1*G85R^C^ mutant, with PA decreasing lifespan in the knock-in mutant strain (**Fig. 3C, S3A-B, Table S7**). BA, 4GBA, 4HP, and PA did not cause consistently significant effects on health span parameters between day 2 to 20 in these strains (**Fig. 3D, S3A-B**). To assess DMD, *dys-1*(*eg33*) and *dys-1*(*cx18*) mutant DMD strains were used, with *dys-1*(*eg33)* demonstrating impaired mitochondrial network integrity and function and a more clinically relevant phenotype than *dys-1*(*cx18)*^44^. BA and 4GBA slightly increased mean lifespan in the *dys-1*(*eg33*) background, whereas PA decreased lifespan, and BA also had the best effect on health span parameters as peristaltic speed and mobility idle time. 4GBA and PA has a negative effect on lifespan in the *dys-1*(*cx18*) background, while no effects were observed for health span (**Fig. 3C-D, S3C-D, Table S7**). Since many neurodegenerations are multifactorial, where both genes and stressors play a role in disease initiation and pathology, we stressed the ALS sod-1WTC and sod-1G85RC mutant line with paraquat and simultaneously supplemented to identify protection by the metabolite candidates. In line with previous studies, healthspan and lifespan decreased after paraquat exposure which was more prominent for the knock-in G85R model (**Fig. 3E-F, Table S8**)^43^. Metabolite supplementation did not induce survival under disease (sod-1 G85R) and paraquat stress, 4HP and 4GBA decreased healthspan and lifespan under disease and paraquat stress (**Fig. 3G-H, Table S8**).

**Figure. 3.**
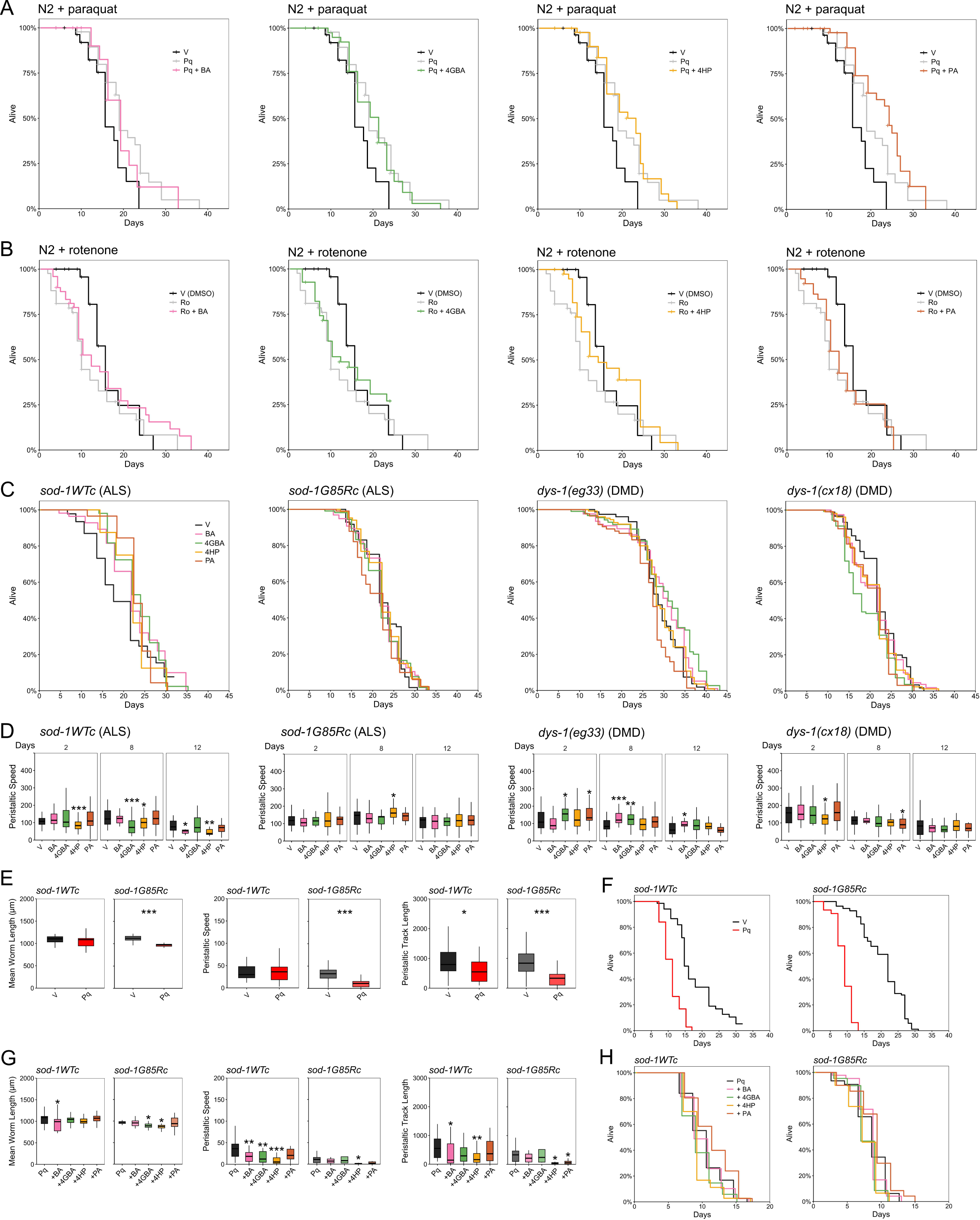
Metabolite candidates overcome oxidative and disease stress in *C. elegans.* (**A, B**) *C. elegans*, wild-type N2, (**C, D**) *C. elegans*, ALS + DMD models (strains HA2986, HA3299, BZ33, LS292). (**A**) Low dose oxidative stress induced by paraquat extended lifespan, which was aggravated by PA, but not by the other metabolite candidates. **(B**) Oxidative stress induced by rotenone was overcome by metabolite candidates; BA, 4GBA and 4HP overcame rotenone stress, (p-values are listed in Table S6). (**C**) Metabolite supplementation did not increase lifespan in an ALS WT control strain, and PA decreased lifespan in an ALS single knock-in strain. 4GBA supplementation increased lifespan in the DMD model (*eg33*), but decreased lifespan in (*cx18*), as did PA supplementation, (p-values are listed in Table S7). (**D**) Health span analysis by peristaltic movement speed identified BA as a health span modulator in the DMD model (*eg33*). No consistent effect by metabolite candidates was observed for the other disease models (ALS and DMD (*cx18*)). (**E**) Health span analysis at day 5 of adulthood by worm length, peristaltic speed and peristaltic track length demonstrated decreased healthspan for paraquat treated sod-1G85R. (**F**) Lifespan analysis demonstrated reduced lifespan of paraquat treated sod-1WTc and sod-1G85R compared to vehicle condition. (**G**) Health span analysis at day 5 of adulthood by worm length, peristaltic speed and peristaltic track length demonstrated decreased healthspan for paraquat and 4HP treated sod-1WTc and sod-1G85R, none of the metabolite candidates saved paraquat induced decreased healthspan. (**H**) Lifespan analysis demonstrated decreased lifespan for paraquat and 4HP and 4GBA treated sod-1WTc and sod-1G85R, none of the metabolite candidates saved paraquat induced decreased lifespan. V: vehicle, BA: β-alanine, 4GBA: 4-guanidinobutanoic acid, 4HP: 4-hydroxyproline, PA: pantothenic acid*. p = * < 0.05, ** < 0.01, *** < 0.001, **** < 0.0001*.

In conclusion, we observed a significant decline of mouse skeletal muscle during aging, which was reflected by an altered skeletal muscle metabolome at an old age. Significantly altered low molecular weight metabolites can impact aspects of longevity and mitochondrial health when supplemented in *C. elegans.* While all the final metabolite candidates (BA, 4GBA, 4HP, PA) affected lifespan, none of them impacted all the parameters studied: lifespan, health span, mitochondrial-, and myosin states, oxidative- and disease stress. This data suggests that a metabolite candidate can show significant effects in one experiment but not in the other. This shows that a successful muscle metabolome screen isolated specific compounds for (only) specific phenotypic and biological aspects of aging. Thus, our work suggests that aging (life and health span) can molecularly be dissected with the metabolite candidates, as these target specific aspects and leave the others untouched. Consequently, our approach can serve as a template and tool for future metabolome screens in healthy or diseased conditions.

## Methods

### Mouse strains, breeding and husbandry

All animal procedures conformed to the National Research Council’s Guide for the Care and Use of Laboratory Animals and were approved by the SingHealth Institutional Animal Care and Use Committee (2019/SHS/1534). C57BL/6J mice were purchased from InVivos Pte Ltd, Singapore, and housed at the SingHealth Experimental Medicine Centre (Academia, Singapore) in a temperature-(22 ± 1 °C), light-(12 h light, 12 h dark) and humidity-controlled (30–40%) environment with free access to food and water. Male and female C57BL/6J mice were aged to young adult (4 months, n=5) and old age (25 months, n=5) after which they were euthanized by cervical dislocation.

### Histology: skeletal muscle

After euthanizing mice, the gastrocnemius and quadriceps were immediately dissected in ice cold HBSS, wiped dry, weighed, and frozen on dry ice. Tissue was stored at –80°C and shipped, kept in dry ice, to Karolinska Institutet for histological sample processing and to the Swedish Metabolomics Centre for metabolomic sample processing. 3 µM muscle sections were prepared with a vibratome, followed by embedding in paraffin wax. Sections were placed on glass slides and baked at 60°C for 1 hour. Immunohistochemistry (IHC) was performed as previously described^45^. Briefly, an automated IHC machine (Leica) was used to perform chromogenic IHC. To avoid batch effects, samples labelled with the same antibody were processed as one batch (list of antibodies in **Table S1**). Wax sections were deparaffinized, dehydrated and antigen was retrieved. Sections were washed and incubated in primary antibody, followed by washing and incubation in a polymer conjugated secondary antibody and colour development. Sections were dehydrated, cleared, mounted, and covered with a glass coverslip.

### Imaging and analysis: skeletal muscle

Whole muscle slices were scanned at 400× with a Grundium Ocus 40 scanner (Grundium Oy, Tampere, Finland). H&E staining was used to analyse total muscle area and muscle fibre area, and to determine five regions of interest (ROIs, 2500 µM^2^). Within the ROIs, fibres were quantified by segmentation using Cell Pose^46^. For fat infiltration quantification, H&E images were manually analysed on adipocyte infiltration by selecting adipocytes cells and -clusters within the whole muscle area. In QuPath, ROIs were analysed on intensity measurements (with background subtraction) for myosin, MHY7B and SDHB. Intensity measurements of single muscle fibres inside the five ROIs were analysed for Tomm20, CyoC, MTCO1, and 8-OHdG^47^.

### Low molecular weight metabolomics: skeletal muscle

Low molecular weight metabolomics was performed as previously described^31^. Briefly, extraction buffer including internal standards were added to the muscle sections together with 1 tungsten bead. Tissues were shaken in a mixer mill and samples were then centrifuged. Supernatant was transferred to micro vials and evaporated to dryness in a speed-vac concentrator. Samples were stored at -80 °C and small aliquots of the remaining supernatants were pooled and used to create quality control (QC) samples. Prior to the analysis, samples were re-suspended in 10 + 10 µL methanol and elution solvent A. The samples were analysed in batches according to a randomized run order. Each batch of samples was first analysed in positive mode. After all samples within a batch had been analysed, the instrument was switched to negative mode and a second injection of each sample was performed. The chromatographic separation was performed on an Agilent 1290 Infinity UHPLC-system (Agilent Technologies, Waldbronn, Germany) and compounds were detected with an Agilent 6546 Q-TOF mass spectrometer equipped with a jet stream electrospray ion source operating in positive or negative ion mode. MSMS analysis was run on the QC samples for identification purposes. All data pre-processing was performed using the Agilent MassHunter Profinder version B.10.0 SP1 (Agilent Technologies Inc., Santa Clara, CA, USA). The quantification of the metabolites was calculated as area under the curve of the mass spectrometry peak and normalized first to an internal standard for negative and positive runs, then for the weight of the tissue. Data were analysed and graphs were made using MetaboAnalyst (version 5.0; 28, 29) and R. All data were subject to Pareto scaling. Hierarchical clustering (HC) (Spearman, Average) was used to create the dendrograms. Comparisons between groups were analysed by two-sample t-tests with an adjusted P value (false discovery rate, FDR), using a cutoff of 0.05 considered significant. Quantitative pathway analysis was performed using the *Mus musculus* KEGG library in MetaboAnalyst 5.0. One animal was excluded from analysis due to outliers.

### Low molecular weight metabolomics: triaging

Metabolites identified in the low molecular weight metabolomics dataset were triaged for follow-up studies in *C. elegans* assays. See **Table 1** for the strategy used.

### C. elegans strains and maintenance

*C. elegans* strains (**Table S2**) and OP50 were obtained from the Caenorhabditis Genetics Centre (CGC, https://cgc.umn.edu/). Strains were maintained on 6 cm dishes at 20°C on nematode growth medium (NGM) with *E. coli* OP50, both prepared according to standard protocols. Chemicals used in *C. elegans* assays are listed in **Table S3**.

### Longevity and health span: C. elegans

All assays were performed on 6 cm dishes and at 20°C. Nematode growth medium (NGM) was prepared according to standard protocols. To prevent the generation of offspring and inhibit egg hatching, 50 µM of FuDR was added to NGM. Compounds were freshly prepared by dissolving in distilled H_2_O or DMSO and sterile filtration. Plates were prepared by supplementing NGM with 0.2 mM, 1 mM or 5 mM of compound (metabolite candidates). Vehicle conditions consisted of 2.5% of distilled H2O or 2.5% of DMSO. Plates were prepared using a peristaltic pump (9 mL) and were dried at RT for 24 hours in the dark.

*E. coli* OP50 as a food source was prepared according to standard protocols, after which it was treated with 1% PFA for 1 hour to metabolically inactivate bacteria^48^. After one hour, OP50 was washed four times with sterile PBS, followed by a 10x concentration in LB and 100 µL per plate was seeded onto NGM plates. Plates were dried at RT for 24 hours in the dark.

A timed egg lay was performed where gravid adults laid eggs for 4 to 5 hours on NGM plates with live OP50. After 4 to 5 hours, gravid adults were removed. After 72 hours, at day 1 of adulthood, L4 to young adult animals were transferred to treatment plates (minimum of 10 and maximum of 30 worms per plate). For flooding assays, at day 12, 16 and 20 of adulthood, 2 mL of M9 buffer was added to the plates and worms were scored as alive (A) when thrashing was visible and scored as dead (D) if worms did not make movements (after poking) and/or were stuck to the plate. Missing worms and worms showing signs of internal hatching were censored (C). For survival assays, worms were scored for live, dead, censored status every 2-4 days. Worms were scored as alive (A) when a reaction to a worm pick was visible and scored as dead (D) if worms did not respond to a worm pick. Missing worms and worms showing signs of internal hatching were censored (C). A log rank test was performed to calculate survival curves and differences in longevity. For health span assays, worms were tracked in real-time on seeded NGM plates by the WormLab system (MBF Biosciences). Plates were gently tapped to stimulate movement and 30 second videos (14 frames per second) were recorded of a selection of worms.

### Chemical and oxidative stress: C. elegans

NGM plates were prepared as described above, paraquat (16 mM) and rotenone (2 µM) were dissolved in 1% PFA OP50 and seeded on 5 mM supplemented plates. For chemical stress assays with wild-type N2, worms were transferred to (fresh) stress plates at day 1 of adulthood and followed up until death. For chemical stress assays with ALS sod-1 worms (sod-1WTc and sod-1G85R), worms were transferred to fresh plates at day 1, 3 and 5 (metabolite candidates were dissolved in NGM and paraquat in OP50 treated with 1% PFA) and paraquat in 100 µL dead OP50 was added every two days from day 7 till death. Worms were scored for live, dead, or censored status every 1-2 days as described above.

### Imaging and analysis: C. elegans

*myo-3p::GFP* and *myo-3p::mitoGFP* worms were anesthetized with 100 mM levamisole hydrochloride and placed on a 2% agarose pad on a glass slide covered by a coverslip. *myo-3p::GFP* worms were imaged with a Leica DMi8, and m*yo-3p::mitoGFP* worms with a Leica Stellaris 5X. For both microscopes, images were acquired with a 63x oil objective with a z-stack of 0.27/0.3 µM respectively. *myo-3p::GFP* images were analysed using the MuscleMetrics plugin in FIJI^39^. *myo-3p::mitoGFP* images were loaded into Imaris (Oxford Instruments), where surfaces were reconstructed. In addition to Imaris, surfaces were reconstructed via the MitoAnalyzer plugin in FIJI^49^.

### Data analysis and statistics

Data distribution was tested by a Shapiro-Wilkinson test and equality of variances was tested by a Levene test. According to the distribution, a T-test or Wilcoxon rank test was performed to compare groups. For metabolomics, a false discovery rate (FDR) of 0.5 and a T-test threshold of 0.1 was set. *C. elegans* lifespan data was analysed by Kaplan-Meier survival curves, p-values were calculated using Log-rank or Cox-regression tests, data was right censored with a death event scored as 1, and lost or censored worms scored as 0. Power analyses were performed to predetermine sample sizes for *C. elegans* microscopy. All statistical analysis and graphics were performed in R.

### Data availability

All data generated is available within the manuscript and its associated supplementary files. Associated protocols and code are available upon request.

## Supplementary Materials

The following supplementary materials are attached:

Supplementary Tables 1-8

Supplementary Figures 1-3

Supplementary Data 1

## Acknowledgments and Funding

The Authors would like to thank the Swedish Metabolomics Centre for help with the metabolic studies, Christian Riedel for advice on *C. elegans* work, and the staff of the histology section at St Erik Eye Hospital. Peter Swoboda was supported by grants from Vetenskapsrådet (2022-04418), the Swedish Brain Foundation (FO2023-0448), and Åhléns Stiftelsen. Pete Williams is supported by St. Erik Eye Hospital’s philanthropic donations, Vetenskapsrådet (2022-00799), a Karolinska Institutet Doctoral education grant, Petrus och Augusta Hedlunds Stiftelse, Ögonfonden, Stiftelsen Kronprinsessan Margaretas Arbetsnämnd, and Karolinska Institutet Foundation grants. *C. elegans* strains were provided by the CGC, which is funded by NIH Office of Research Infrastructure Programs (P40 OD010440).

## Author contributions

SMJ – performed experiments and analysis, conceived and designed experiments, wrote the manuscript. AN – performed experiments. VC – performed experiments. EL – performed experiments. SY – performed experiments. GS – performed experiments, provided supervision and expertise. JGC – provided supervision and expertise. JRT – performed experiments, provided supervision. PS – provided supervision and expertise. PAW – provided supervision, conceived and designed experiments, wrote the manuscript. All authors read and approved of the final manuscript.

---

**Figure. S1.**
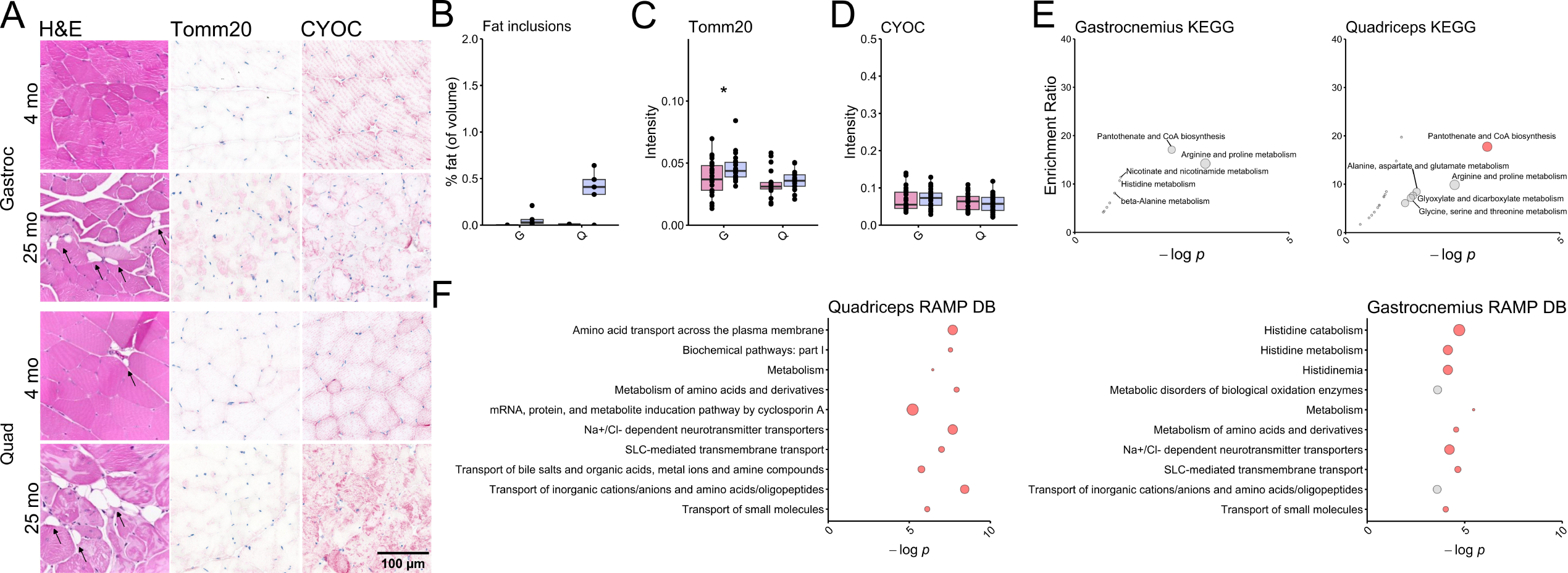
Additional skeletal muscle: histology and metabolomics. (**A-F**) C57BL/6J mice. (**A**) By histological assessment, it was observed that (**B**) there was a slight increase of adipocyte infiltration during aging in the quadriceps muscle (ns), (**C**) Tomm20 intensity increased, but (**D**) CYOC intensity did not change. (**E**) A KEGG enrichment analysis shows the top 5 affected pathways in gastrocnemius and quadriceps, of which only pantothenate and CoA biosynthesis were significant. (**F**) Ramp DB enrichment shows the top 10 processes that significant hits are involved in. For KEGG and Ramp DB plots, red dots show an FDR<0.05, grey dots show an FDR>0.05 and dot size is based on the number of hits (between 1-3) and the enrichment ratio (between 3-13). *Pink* = 4 months of age, *purple* = 25 months of age. *p = * < 0.05, ** < 0.01, *** < 0.001, **** < 0.0001*.

**Figure. S2.**
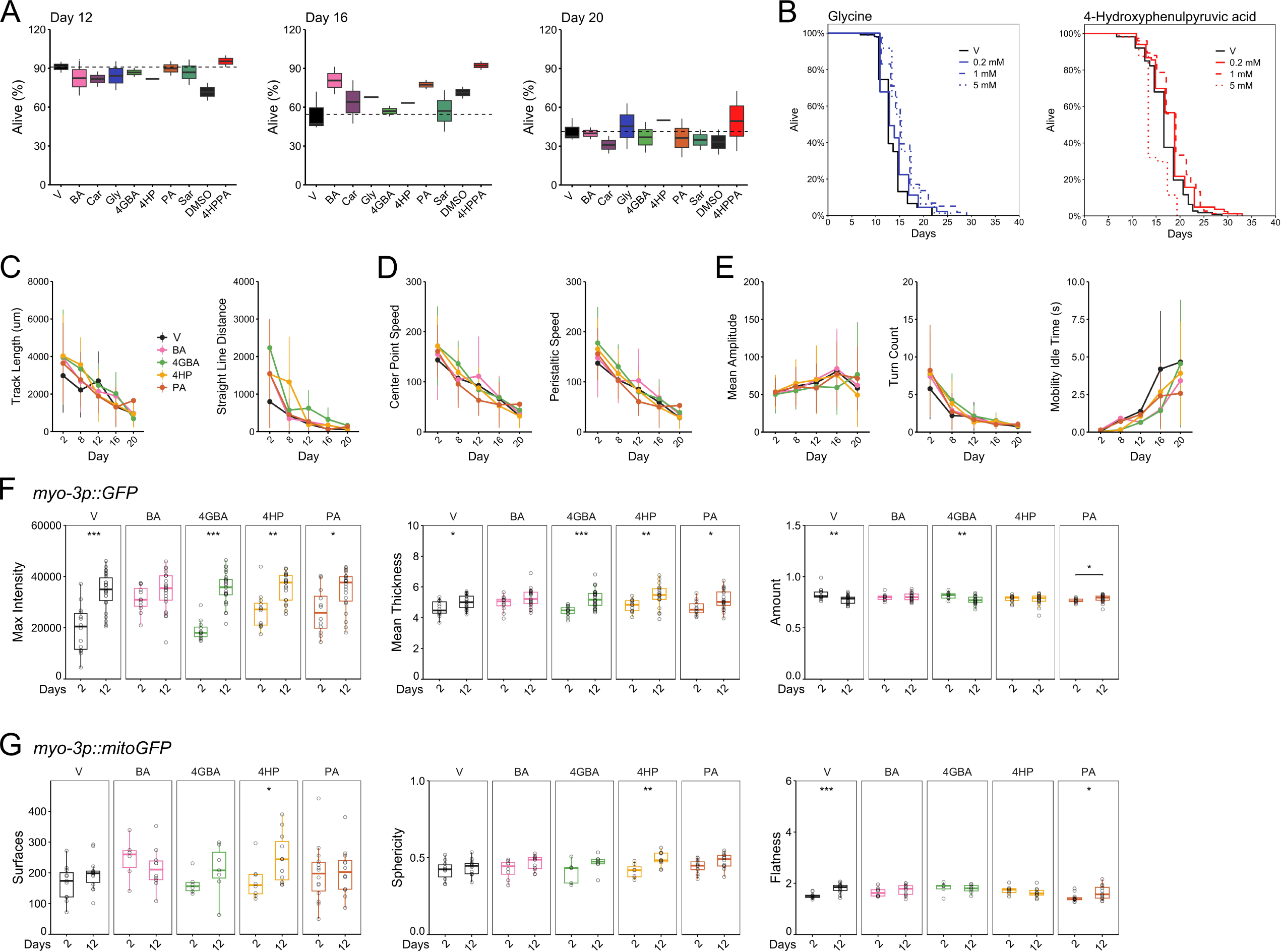
Additional *C. elegans*: longevity, WormLab and microscopy. (**A-E**) *C. elegans,* wild-type N2, (**F**) *C. elegans*, RW1596, (**G**) *C. elegans*, SJ4103. (**A**) Flooding assays identify potential health span enhancers at day 16 and lifespan enhancers at day 20. Based on these results BA, 4HP, 4GBA and PA were selected for survival analysis. Survival analysis after (**B**) glycine supplementation showed a slight increase of lifespan, and 4HPPA supplementation showed a slight increase at 0.2 and 1 mM, but a toxic effect at 5 mM. However, the compound was dissolved in DMSO, which by itself increased lifespan compared to wild-type N2 and vehicle (ddH_2_O). Based on survival curves, BA, 4GBA, 4HP and PA were selected for health span follow up (**Table S5**), identifying slight increases at D2 of adulthood for movement (**C**) distance parameters, and (**D**) speed parameters. (**E**) Amplitude and turn count were similar for all conditions, while mobility idle time was highest for the vehicle condition later in life (day 16 and 20). (**F**) Fluorescence microscopy images of *myo-3p::GFP* (RW1596) were quantified, where an increase of max intensity is identified due to the intensity of the blebs, the mean thickness increased, while the amount decreased. BA did not show any significant effects, suggesting a protective effect on myosin during aging. (**G**) Fluorescence confocal microscopy images of *myo-3p::mitoGFP* (SJ4103) were quantified, where a similar number of mitochondrial surfaces was identified as well as a similar sphericity. Flatness increased in the vehicle and PA condition. V: vehicle, BA: β-alanine, Car: carnosine, Gly: glycine, 4GBA: 4-guanidinobutanoic acid, 4HP: 4-hydroxyproline, PA: pantothenic acid, Sar: sarcosine, DMSO: dimethyl sulfoxide, 4HPPA: 4-hydroxyphenylpyruvic acid. *p = * < 0.05, ** < 0.01, *** < 0.001, **** < 0.0001*.

**Figure. S3.**
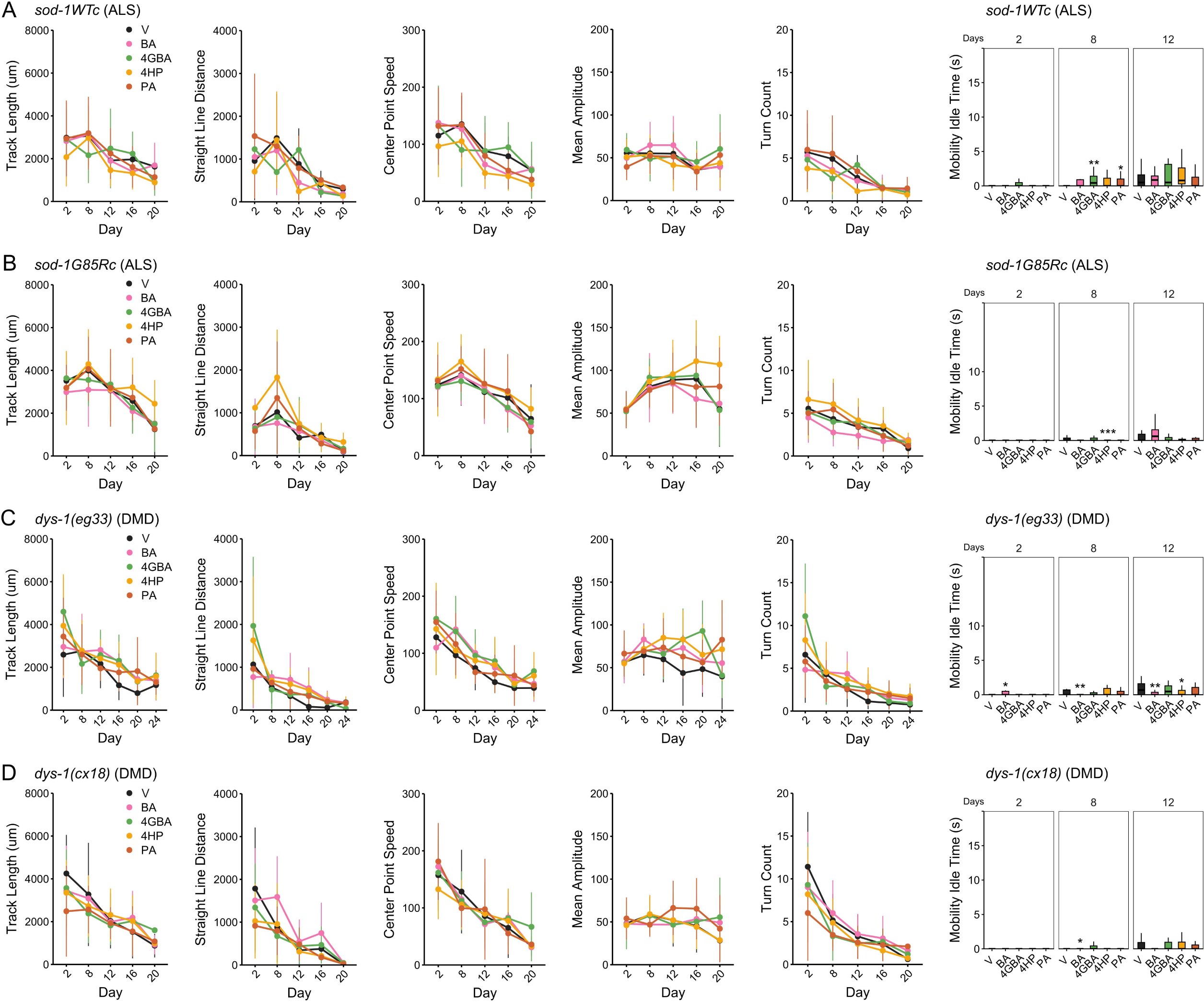
Additional *C. elegans:* oxidative and genetic stress. *C. elegans* strains: (**A**) HA2986, (**B**) HA3299, (**C**) BZ33, (**D**) LS292. All panels show health span parameters assessed by WormLab. (**A**) For the ALS wild-type control (*sod-1*WTc) strain, all conditions show overlap in different health span parameters, which is also observed for the (**B**) ALS knock-in (*sod-1*G85Rc) strain. (**C**) For the DMD (*dys-1*(*eg33*)) strain all metabolite candidates appear to increase performance on health span parameters compared to the vehicle control condition, while this is not observed for the (**D**) DMD (*dys-1*(cx18)) strain. V: vehicle, BA: β-alanine, 4GBA: 4-guanidinobutanoic acid, 4HP: 4-hydroxyproline, PA: pantothenic acid. *p = * < 0.05, ** < 0.01, *** < 0.001, **** < 0.0001*.

**Supplementary Table 1.** Antibodies used.

| Target | Clone | # | Supplier | Dilution |
| --- | --- | --- | --- | --- |
| CYOC | Rabbit | ab150422 | Abcam | 0.32 µg/µl (1:700) |
| MTCO1 | Mouse | ab14705 | Abcam | 1:200 |
| MYH7B | Rabbit | ab240173 | Abcam | 10.46 µg/µl (1:100) |
| Myosin | Mouse | ab37484 | Abcam | 0.81-0.99 µg/µl (1:1000) |
| SDHB | Mouse | ab14714 | Abcam | 1:100 |
| Tomm20 | Rabbit | ab78547 | Abcam | 0.9-1 µg/µl (1:1000) |
| 8-OHdG | Rabbit | bs-1278R | Bioss Biosciences | 0.01 µg/µl (1:100) |

**Supplementary Table 2.** List of *C. elegans* strains.

| Strain name | Genotype | Relevant details |
| --- | --- | --- |
| N2 | Wild type strain. |  |
| BZ33 | *dys-1(eg33) I.* | *dys-1*: dystrophin related<br>(*eg33*): substitution allele |
| HA2986 | *sod-1(rt448[sod-1WTc]) II.* | *sod-1*: superoxide dismutase<br>WTc: CRISPR/Cas9 with silent codon change |
| HA3299 | *sod-1(rt451[sod-1G85R]) II.* | *sod-1*: superoxide dismutase<br>1G85R: CRISPR/Cas9 G85R missense mutation in *sod-1* |
| LS292 | *dys-1(cx18) I.* | *dys-1*: dystrophin related<br>(*cx18*): allele |
| SJ4103 | *zcls14 [myo-3::GFP(mit)]* | *zcls14*: [*myo-3::gfp(mit)*] |
| RW1596 | *myo-3(st386) V; stEx30.* | *myo-3*: myosin heavy chain<br>*st386*: allele<br>*stEx30*: [*myo-3p::GFP::myo-3 + rol-6*] |

**Supplementary Table 3.**
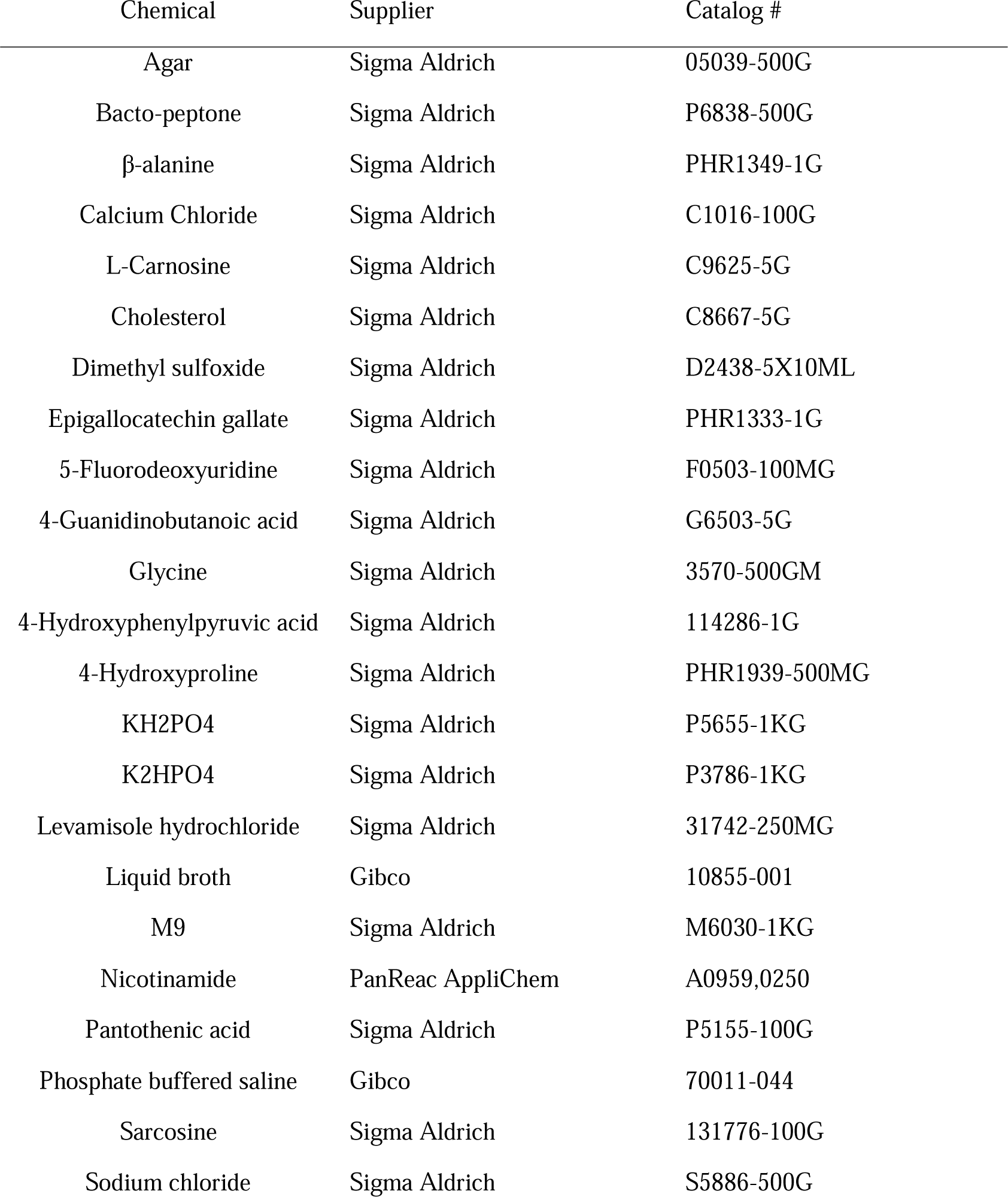
List of chemicals used in *C. elegans* assays.

| Chemical | Supplier | Catalog # |
| --- | --- | --- |
| Agar | Sigma Aldrich | 05039-500G |
| Bacto-peptone | Sigma Aldrich | P6838-500G |
| β-alanine | Sigma Aldrich | PHR1349-1G |
| Calcium Chloride | Sigma Aldrich | C1016-100G |
| L-Carnosine | Sigma Aldrich | C9625-5G |
| Cholesterol | Sigma Aldrich | C8667-5G |
| Dimethyl sulfoxide | Sigma Aldrich | D2438-5X10ML |
| Epigallocatechin gallate | Sigma Aldrich | PHR1333-1G |
| 5-Fluorodeoxyuridine | Sigma Aldrich | F0503-100MG |
| 4-Guanidinobutanoic acid | Sigma Aldrich | G6503-5G |
| Glycine | Sigma Aldrich | 3570-500GM |
| 4-Hydroxyphenylpyruvic acid | Sigma Aldrich | 114286-1G |
| 4-Hydroxyproline | Sigma Aldrich | PHR1939-500MG |
| KH<sub>2</sub>PO<sub>4</sub> | Sigma Aldrich | P5655-1KG |
| K<sub>2</sub>HPO<sub>4</sub> | Sigma Aldrich | P3786-1KG |
| Levamisole hydrochloride | Sigma Aldrich | 31742-250MG |
| Liquid broth | Gibco | 10855-001 |
| M9 | Sigma Aldrich | M6030-1KG |
| Nicotinamide | PanReac AppliChem | A0959,0250 |
| Pantothenic acid | Sigma Aldrich | P5155-100G |
| Phosphate buffered saline | Gibco | 70011-044 |
| Sarcosine | Sigma Aldrich | 131776-100G |
| Sodium chloride | Sigma Aldrich | S5886-500G |

**Supplementary Table 4.** Summary results of *C. elegans* flooding assays.

| Compound | Dose (mM) | Day 16 (% alive, ±, n) | Day 20 (% alive, ±, n) |
| --- | --- | --- | --- |
| Vehicle (ddH<sub>2</sub>O) | 2.5% | 55 ± 15, n=96 | 41 ± 9, n=101 |
| β-Alanine | 5 | 81 ± 18, n=73 | 40 ± 6, n=61 |
| Carnosine | 5 | 64 ± 24, n=64 | 31 ± 10, n=70 |
| Glycine | 5 | 68 ± 1, n=77 | 45 ± 25, n=74 |
| 4-Guanidinobutanoic acid | 5 | 57 ± 6, n=53 | 37 ± 17, n=67 |
| 4-Hydroxyproline | 5 | 63, n=49 | 50, n=44 |
| Pantothenic acid | 5 | 77 ± 5, n=63 | 36 ± 21, n=71 |
| Sarcosine | 5 | 57 ± 23, n=60 | 35 ± 11, n=91 |
| DMSO | 2.5% | 71 ± 7, n=81 | 33 ± 14, n=59 |
| 4-Hydroxyphenyl pyruvic acid | 5 | 92 ± 5, n=40 | 49 ± 33, n=34 |

\*n=number of events (censored worms excl.)

**Supplementary Table 5.**
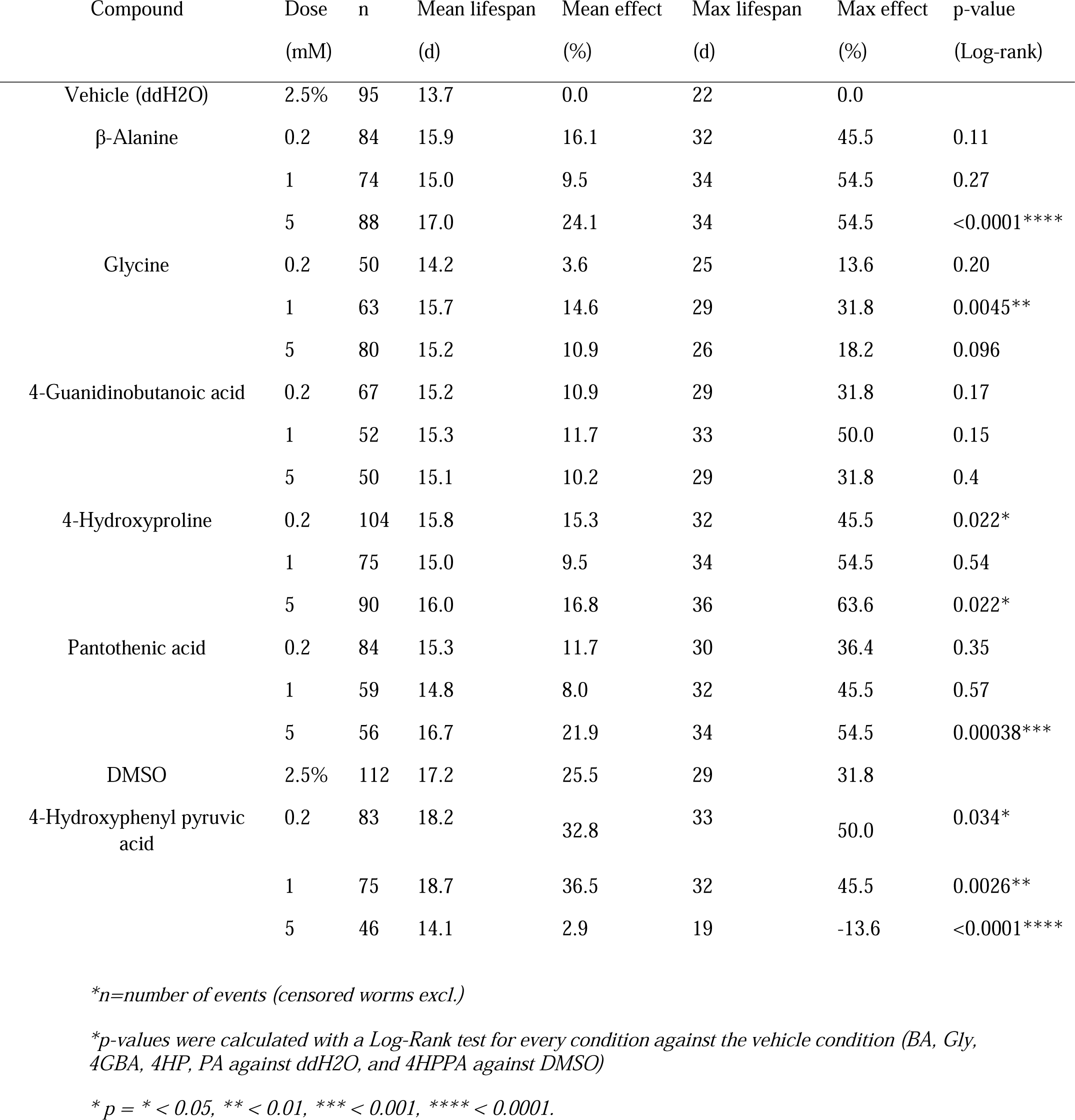
Summary results of *C. elegans* survival assay.

**Supplementary Table 6.**
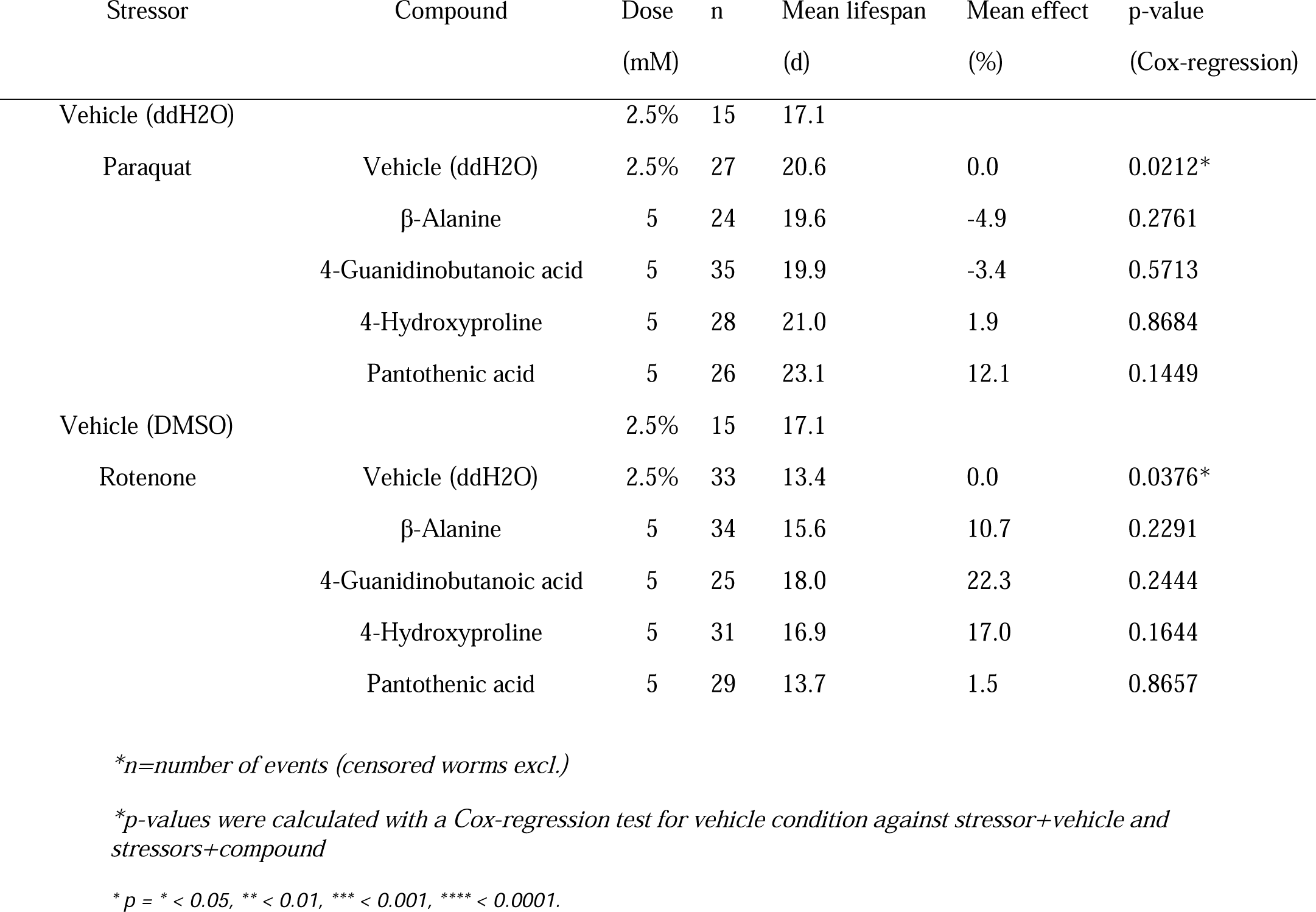
Summary results of *C. elegans* oxidative stress survival assay.

**Supplementary Table 7.** Summary results of mutant *C. elegans* strains survival assay.

| Mutant strain<br>(genotype) | Compound | Dose<br>(mM) | n | Mean lifespan<br>(d) | Mean effect<br>(%) | p-value<br>(Cox-regression) |
| --- | --- | --- | --- | --- | --- | --- |
| HA2986 | Vehicle (ddH<sub>2</sub>O) | 2.5% | 33 | 20.2 | 0.0 |  |
| *(sod-1(rt448[sod-1WTc]) II)* | β-Alanine | 5 | 51 | 22.7 | 12.4 | 0.0787 |
|  | 4-Guanidinobutanoic acid | 5 | 42 | 23.4 | 15.8 | 0.1174 |
|  | 4-Hydroxyproline | 5 | 8 | 22.0 | 8.9 | 0.8740 |
|  | Pantothenic acid | 5 | 24 | 22.7 | 12.4 | 0.5500 |
| HA3299 | Vehicle (ddH<sub>2</sub>O) | 2.5% | 73 | 22.9 | 0.0 |  |
| *(sod-1(rt451[sod-1G85R]) II)* | β-Alanine | 5 | 67 | 22.5 | -1.7 | 0.9377 |
|  | 4-Guanidinobutanoic acid | 5 | 85 | 22.1 | -3.5 | 0.6058 |
|  | 4-Hydroxyproline | 5 | 73 | 22.5 | -1.7 | 0.6971 |
|  | Pantothenic acid | 5 | 85 | 20.7 | -9.6 | 0.0336* |
| BZ33 | Vehicle (ddH<sub>2</sub>O) | 2.5% | 68 | 29.4 | 0.0 |  |
| *(dys-1(eg33) I)* | β-Alanine | 5 | 109 | 29.8 | 1.4 | 0.118899 |
|  | 4-Guanidinobutanoic acid | 5 | 76 | 30.5 | 3.7 | 0.00650** |
|  | 4-Hydroxyproline | 5 | 115 | 28.8 | -2.0 | 0.96836 |
|  | Pantothenic acid | 5 | 77 | 26.3 | -10.5 | 0.00184** |
| LS292 | Vehicle (ddH<sub>2</sub>O) | 2.5% | 75 | 22.7 | 0.0 |  |
| *(dys-1(cx18) I)* | β-Alanine | 5 | 111 | 21.4 | -5.7 | 0.281233 |
|  | 4-Guanidinobutanoic acid | 5 | 101 | 19.1 | -15.9 | 0.000308*** |
|  | 4-Hydroxyproline | 5 | 90 | 20.8 | -8.4 | 0.052219 |
|  | Pantothenic acid | 5 | 69 | 20.0 | -11.9 | 0.017031* |

\*n=number of events (censored worms excl.)
\*p-values were calculated with a Cox-regression test for vehicle condition against compound per genetic background
\* $p = * < 0.05$ , \*\* $< 0.01$ , \*\*\* $< 0.001$ , \*\*\*\* $< 0.0001$ .

**Supplementary Table 8.** Summary results of mutant sod-1 *C. elegans* strains under oxidative stress.

| Mutant strain<br>(genotype) | Stressor | Compound | Dose<br>(mM) | n | Mean lifespan<br>(d) | p-value<br>(Cox-regression) |
| --- | --- | --- | --- | --- | --- | --- |
| HA2986 | Vehicle (ddH<sub>2</sub>O) | Vehicle (ddH<sub>2</sub>O) | 2.5% | 50 | 18.1 |  |
| *(sod-1(rt448[sod-1WTc]) II)* | Paraquat (20 mM) | Vehicle (ddH<sub>2</sub>O) | 5 | 41 | 10.7 | 1.37E-12**** |
|  |  | β-Alanine | 5 | 41 | 10.4 | 0.59615 |
|  |  | 4-Guanidinobutanoic acid | 5 | 39 | 9.5 | 0.04948* |
|  |  | 4-Hydroxyproline | 5 | 37 | 9.1 | 0.00838** |
|  |  | Pantothenic acid | 5 | 39 | 11.1 | 0.55291 |
| HA3299 | Vehicle (ddH<sub>2</sub>O) | Vehicle (ddH<sub>2</sub>O) | 2.5% | 96 | 20.7 |  |
| *(sod-1(rt451[sod-1G85R]) II)* | Paraquat (20 mM) | Vehicle (ddH<sub>2</sub>O) | 5 | 33 | 8.8 | <2E-16**** |
|  |  | β-Alanine | 5 | 25 | 8.7 | 0.61266 |
|  |  | 4-Guanidinobutanoic acid | 5 | 26 | 7.8 | 0.03191* |
|  |  | 4-Hydroxyproline | 5 | 23 | 7.33 | 0.00552** |
|  |  | Pantothenic acid | 5 | 29 | 8.7 | 0.96923 |

\*n=number of events (censored worms excl.)
\*p-values were calculated with a Cox-regression test in vehicle condition for vehicle against paraquat and in paraquat condition for vehicle against compounds
\* $p = * < 0.05$ , \*\* $< 0.01$ , \*\*\* $< 0.001$ , \*\*\*\* $< 0.0001$ .

